# Effects of face masks and ventilation on the risk of SARS-CoV-2 respiratory transmission in public toilets: a quantitative microbial risk assessment

**DOI:** 10.1101/2021.08.21.457245

**Authors:** Thammanitchpol Denpetkul, Oranoot Sittipunsakda, Monchai Pumkaew, Skorn Mongkolsuk, Kwanrawee Sirikanchana

## Abstract

Public toilets could increase the risk of COVID-19 infection via airborne transmission; however, related research is limited. We aimed to estimate SARS-CoV-2 infection risk through respiratory transmission using a quantitative microbial risk assessment framework by retrieving SARS-CoV-2 concentrations from the swab tests of 251 Thai patients. Three virus-generating scenarios were investigated: an infector breathing, breathing with a cough, and breathing with a sneeze. Infection risk (97.5th percentile) was as high as 10^−3^ with breathing and increased to 10^−1^ with a cough or sneeze, thus all higher than the risk benchmark of 5 × 10^−5^ per event. No significant gender differences for toilet users (receptors) were noted. The highest risk scenario of breathing and a sneeze was further evaluated for risk mitigation measures. Risk mitigation to lower than the benchmark succeeded only when the infector and receptor simultaneously wore an N95 respirator or surgical mask and when the receptor wore an N95 respirator and the infector wore a denim fabric mask. Ventilation up to 20 air changes per hour (ACH), beyond the 12-ACH suggested by the WHO, did not mitigate risk. Virus concentration, volume of expelled droplets, and receptor dwell time were identified as the main contributors to transmission risk.

**Highlights:**

- The use of public toilets poses a risk of SARS-CoV-2 respiratory transmission
- Highest risks generated in the order of sneezing, coughing, and breathing
- No gender differences in risk by counteracting dwell times and inhalation rates
- Ventilation did not reduce risk even at 20 ACH, beyond the WHO-recommended value
- N95 and surgical masks offer the most effective risk mitigation to toilet users

**Graphical abstract:** 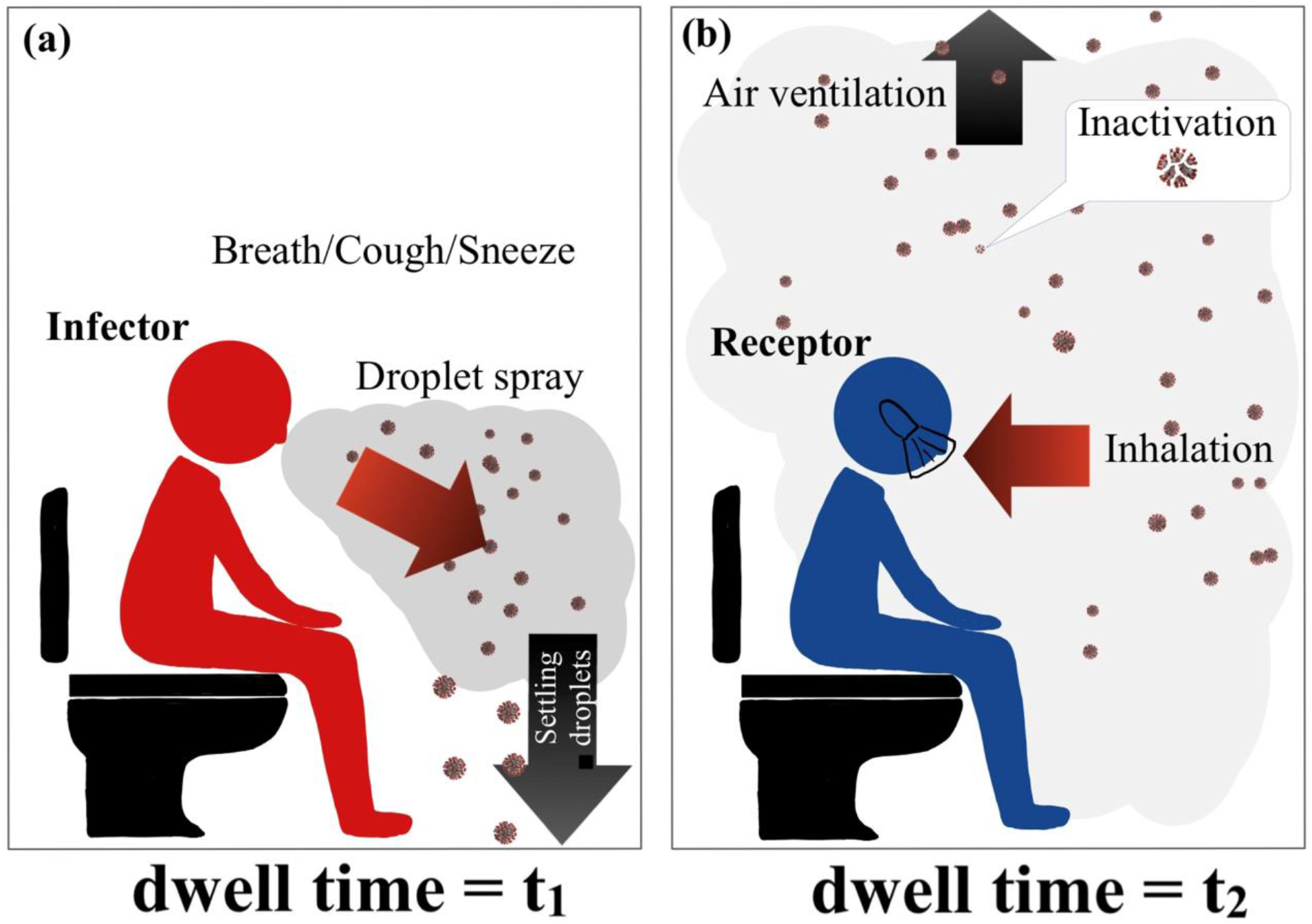

## Introduction

The coronavirus disease 2019 (COVID-19) pandemic has affected the global population since its first emergence in December 2019. The three main transmission routes of the severe acute respiratory syndrome coronavirus 2 (SARS-CoV-2), an etiological agent of COVID-19, have been identified as (1) the inhalation of respiratory fluids carrying infectious viruses, (2) direct splashes or sprays of infectious respiratory droplets and aerosol particles, and (3) the touching of contaminated surfaces (Centers for Disease Control and Prevention, 2021). Communal confined spaces such as public toilets in shopping centers, schools, restaurants, airports, theaters, and hospitals may be significant areas for SARS-CoV-2 transmission (Dancer, Li, et al., 2021). Surface contamination with SARS-CoV-2 in toilets and bathrooms has been reported (Ding, Qian, et al., 2021; Maestre, Jarma, et al., 2021); however, transmission risks from fomite exposure could be reduced significantly through the simple yet effective interventions of hand washing, hand sanitizing, and surface disinfection (Dancer et al., 2021; World Health Organization [WHO], 2020a; Pitol & Julian, 2021). Airborne transmission, on the other hand, is deemed the main route of COVID-19 spread (Centers for Disease Control and Prevention, 2021) and could be aggravated by the use of busy, confined public toilet spaces, especially if appropriate steps are not taken to mitigate the risk of virus transmission (Dancer, Li, et al., 2021).

The positive detection of SARS-CoV-2 RNA has been reported in 23.8% of air samples from hospital toilets, which have demonstrated higher viral loads than clinical areas (Birgand, Peiffer-Smadja, et al., 2020). However, risk assessments of respiratory exposure to SARS-CoV-2 in public toilets is limited. Potential sources of infectious respiratory droplets and aerosol particles in toilet settings include exhalation and expelling, such as sneezing, speaking, and coughing, by infected toilet users, and the aerosolization of infected feces and urine after toilet flushing (Dancer, Li, et al., 2021; Schijven, Vermeulen, et al., 2021). Although infectious SARS-CoV-2 was isolated from the feces of a severely infected patient (Xiao, Sun, et al., 2020), studies confirming that feces and urine in wastewater remain infectious for SARS-CoV-2 are limited, with supporting evidence showing poor virus survival in gastrointestinal tracts due to the low pH of gastric fluids, bile, digestive enzymes, and bacterial byproducts (Zang, Castro, et al., 2020; Albert, Ruíz, et al., 2021; Jones, Baluja, et al., 2020). Consequently, even though flushing activities can produce airborne droplets and aerosols, the associated risks may be low because contamination by infectious virus particles is less likely (Shi, Huang, et al., 2021). In this study, we therefore focused on characterizing the risk of SARS-CoV-2 respiratory transmission introduced by normal breathing and expelling (i.e., coughing and sneezing) in a public toilet setting.

Quantitative microbial risk assessment (QMRA) is a valuable tool used to quantitatively estimate human health risks associated with exposure to pathogens in different environmental matrices (Rose and Gerba, 1991; Haas, Rose, et al., 2014). The QMRA framework has been applied to estimate SARS-CoV-2 transmission risk to wastewater treatment plant workers (Dada and Gyawali, 2021; Zaneti, Girardi, et al., 2021), Tokyo 2020 Olympic Games attendees (Murakami, Miura, et al., 2021), and confined vehicle passengers and shared room users (Schijven, Vermeulen, et al., 2021). In the present study, we aimed to estimate the risk of infection associated with public toilet exposure to SARS-CoV-2 through airborne transmission using the QMRA approach. For convenience, we called a healthy person who is exposed to transmission risk a receptor, while a disease-carrying person, either symptomatic or asymptomatic, was termed an infector. We gathered the input parameters from a variety of sources. These included COVID-19 concentration data obtained by swab testing 251 Thai patients from a public hospital in Bangkok and the exposure factors related to three droplet- and aerosol-generating activities of infectors, namely, breathing, breathing with a cough, and breathing with a sneeze, which were all identified from published sources (Schijven, Vermeulen, et al., 2021; Fabian, Brain, et al., 2011; Duguid, 1946; Han, Weng, et al., 2013; Loudon and Roberts, 1967). The risk of infection was calculated separately for male and female receptors because of their different respiratory rates and periods of time spent in the toilet, the so-called dwell times. To include uncertainty and variability in the risk characterization, we applied the Monte Carlo simulation technique to calculate the risks. The sensitivity of the model parameters was evaluated to determine which input parameters could help reduce the associated uncertainty. Finally, two risk mitigation measures, namely, face mask wearing and ventilation improvement, were assessed to ascertain their efficacy. The calculated risks and associated mitigation measures may be beneficial in the development of public health policies aimed at providing effective control of SARS-CoV-2 transmission.

## MATERIALS AND METHODS

### Risk scenarios

We evaluated the infection risk in various scenarios with and without preventive measures. For the public toilet model, a Thailand standard cubicle size of 1.5 × 0.8 × 2.7 m (3.24 m^3^) was set for the risk evaluation (Ministry of Public Health, 2016). The three scenarios used in this study that can cause an infector to generate infectious droplets and aerosols included breathing (Br), breathing with a cough (Br+Co), and breathing with a sneeze (Br+Sn). The scenario that provided the highest risk was further investigated to determine the efficacy of the identified mitigation measures (i.e., face mask wearing and ventilation). To evaluate the effects of mask wearing, different types of masks (i.e., N95 respirator and surgical and denim fabric masks) were modeled when worn by either an infector or a receptor, or both. For the ventilation evaluation, the air changes per hour (ACH) were varied at 0 (no ventilation), 0.5 (poor ventilation), 10 (DIN 1946 ventilation standard for public toilets), 12 (WHO recommended standard ventilation [2021]), and 20 (extreme ventilation). An outline of the QMRA steps for all the scenarios are presented in Figure 1.

**Figure 1.**
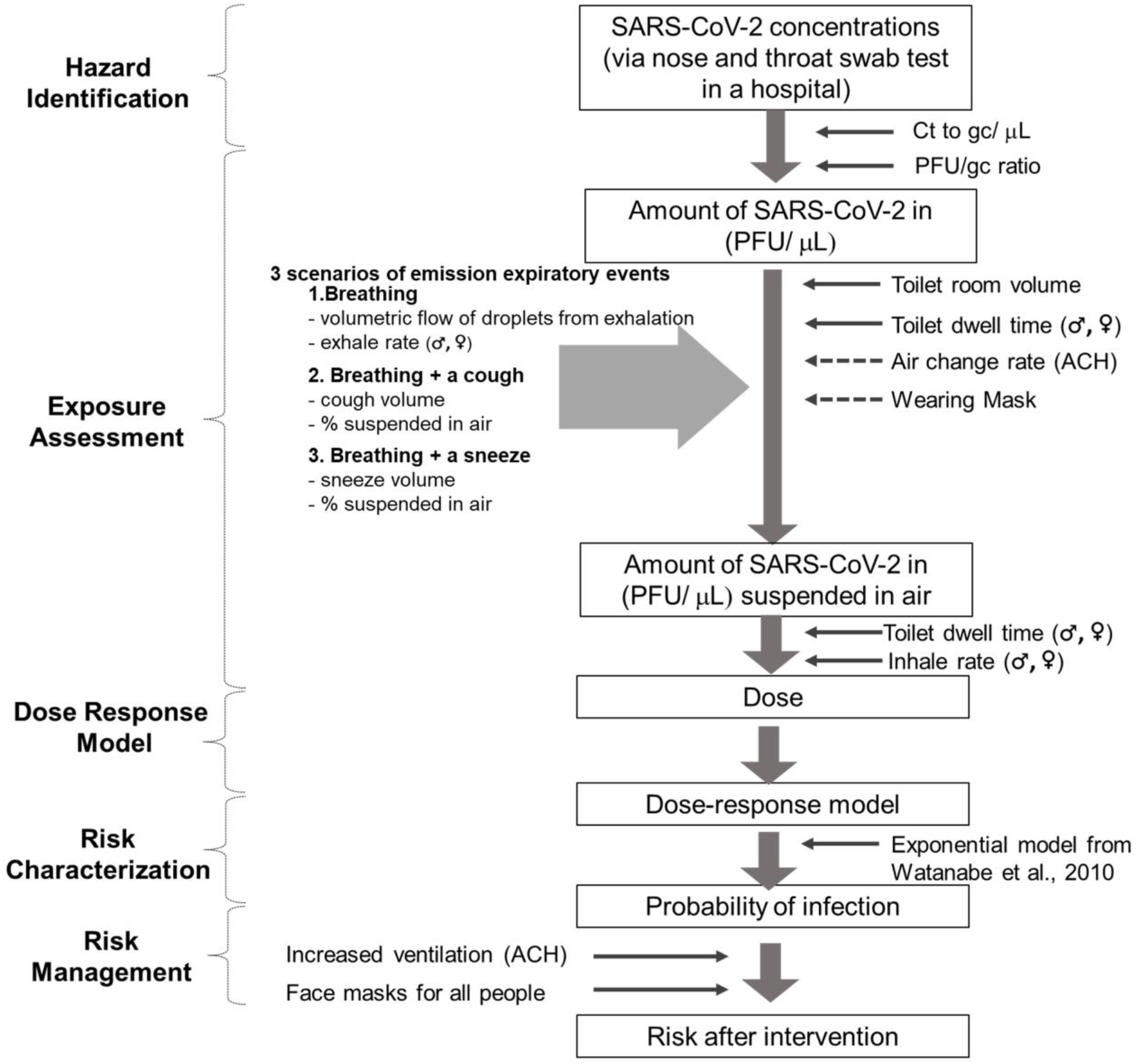
Outline of the QMRA steps for the scenarios associated with SARS-CoV-2 respiratory transmission in a public toilet setting

### Virus levels generated by an infector

The SARS-CoV-2 concentrations used in this study were retrieved from the reverse-transcription quantitative polymerase chain reaction quantification cycle (C_t_) values of 251 positive swab test results of an N2 gene from a public hospital in Bangkok from March to May 2021. Due to the absence of a standard curve for clinical swab testing in Thailand, the viral concentrations were estimated using a published standard curve (Sherchan, Shahin, et al., 2020), which ranged from 4.4 × 10^−1^ to 6.4 × 10^8^ gene copies (gc)/μL. The SARS-CoV-2 concentrations (*A*) were fitted with a triangular distribution as shown in Table S1. To convert the virus concentrations from gc to an infectious plaque-forming unit (PFU), the ratios of the PFU/gc (*R*) of SARS-CoV-2 between 1:100 and 1:1000 with a uniform distribution were applied (Pitol & Julian, 2021).

The number of infectious viruses suspended in the ambient air of the toilet cubicle was calculated using the mass balance equation (Eq. 1) in which each term in the equation has units of mass per time. Under the completely mixed condition, the accumulation of virus particles as aerosols was obtained from the summation of the breathing of an infector, virus inactivation, and virus removal by mechanical ventilation and inhalation. Under typical conditions of 20%–70% relative humidity, a 20°C temperature, and no direct sunlight, an average SARS-CoV-2 inactivation rate of 0.008 (min^−1^) was applied (Schuit, Ratnesar-Shumate, et al., 2020). We assumed that the virus particles were released continually during the time the infector spent in the toilet cubicle (infector’s dwell time = *t_1_* minutes). The dwell times for men and women, which were in line with those indicated in an airport study (250 men and 237 women), were fitted with a log-normal distribution (Table S1) (Gwynne, Hunt, et al., 2019). The remaining infectious virus concentrations generated by the infector after leaving the toilet (*C_t1_*) were calculated according to Eq. 2 by integrating Eq. 1 with no initial virus particles (*C* = 0). The inhalation rates (*q_in_*) following the uniform distribution ranged between 8.36 and 19.74 L/min for men and 6.4 and 13.78 L/min for women (Brochu, Ducré-Robitaille, et al., 2006). The additional concentrations of infectious SARS-CoV-2 expelled by coughing (*C_co_*) and sneezing (*C_sn_*) were calculated using Eqs. 3 and 4, respectively. The volumetric flow of droplets from an infector’s exhalation (*q_br_*) ranged from 5 × 10^−9^ to 6 × 10^−6^ μL/min (Schijven, Vermeulen, et al., 2021). In addition, the volume of aerosol droplets expelled per cough (*V_co_*) and per sneeze (*V_sn_*) was set according to the literature (Schijven, Vermeulen, et al., 2021). However, the size of the droplets played an important role in their activity. The larger droplets deposited quickly, whereas the smaller droplets (aerosols) could remain suspended in the air for a longer period. Thus, the volumetric ratios of aerosols to total droplets expelled (*F*) were considered using a droplet size ≤70 μm based on the size distribution of droplets in the literature for breathing (Fabian, Brain, et al., 2011), coughing (Duguid, 1946; Han, Weng, et al., 2013; Loudon and Roberts, 1967) and sneezing (Duguid, 1946; Han, Weng, et al., 2013) (Table S1).

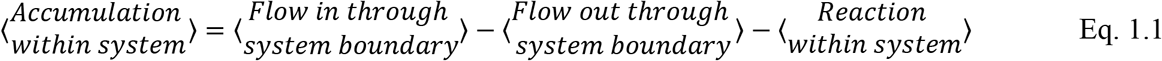

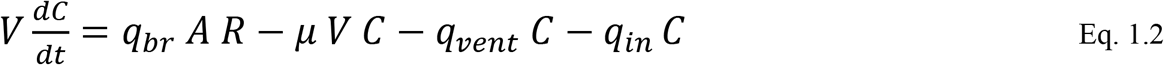

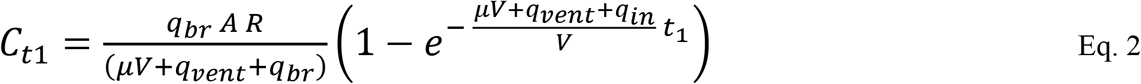

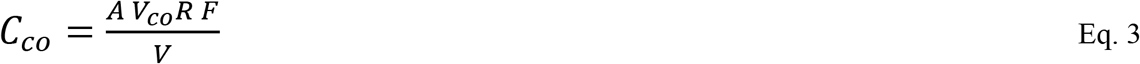

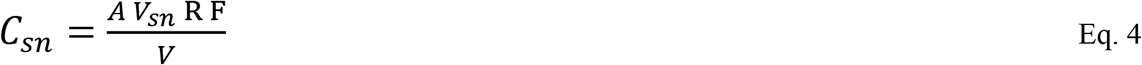

where the parameters related to virus generation by an infector are:

*q_br_* = volumetric flow of droplets from an infector’s exhalation (μL-droplet/min)
*A* = virus concentrations in the genome copies per volume of droplets (gc/μL-droplet)
*C_co_* = additional infectious virus concentrations in the air caused by a cough (PFU/L)
*C_sn_* = additional infectious virus concentrations in the air caused by a sneeze (PFU/L)
*R* = PFU/gc ratio
*F* = fraction of aerosol volume per total volume of droplets expelled (dimensionless)
*V_co_* = volume of aerosol expelled per cough (μL/cough)
*V_sn_* = volume of aerosol expelled per sneeze (μL/sneeze)
*q_in_* = inhalation rate (L/min)
*q_vent_* = ventilation rate (L/min) that equals ACH × V/60
*V* = volume of air in a cubicle (3,240 L-air)
*t_1_* = infector’s dwell time (min)
*μ* = inactivation rate in the air at 20%–70% relative humidity levels (min^−1^)

### Virus levels accessible to a receptor

Because no input source of virus (no infector) was present, the term of flow in through a system boundary was discarded. To solve Eq. 5, the infectious virus concentrations the receptor (*C_t2_*) was exposed to during the receptor’s dwell time *t_2_* were calculated from Eq. 6. In this study, we assumed that after the infector exited the cubicle, the receptor immediately entered the cubicle. The initial concentrations (*C_0_*) based on three scenarios (i.e., breathing only, breathing with a cough, and breathing with a sneeze [Eqs. 7–9]) were therefore incorporated into Eq. 6.

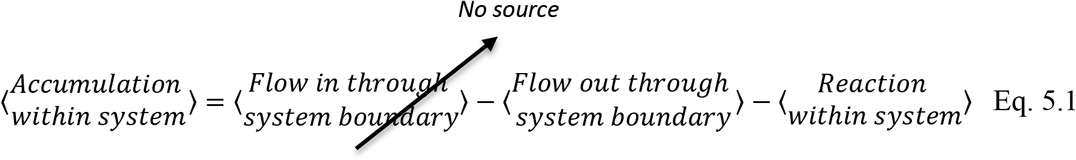

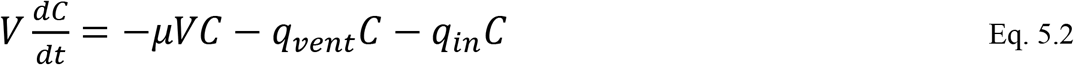

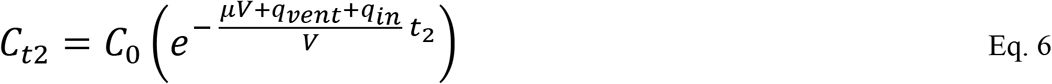

Given *C_0_* based on the following specific scenarios:

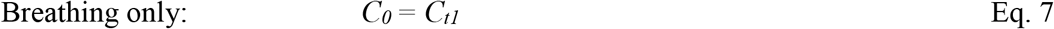

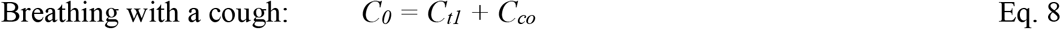

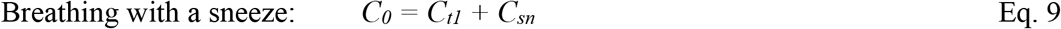

### Virus doses inhaled by the receptor

The virus doses (*d*) that would be inhaled by a receptor were calculated by incorporating the inhalation rate (*q_in_*) with a definite integral of the infectious virus concentration-time function (*C_t2_*). The limits of integration were set from *t* = 0 to the receptor’s dwell time (*t_2_*) (Eqs. 10.1–10.2). The initial concentrations (*C_0_*) also followed Eqs. 7–9 in line with the desired scenarios.

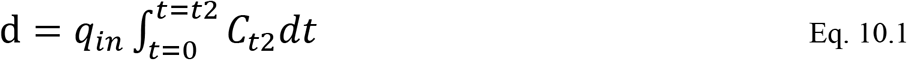

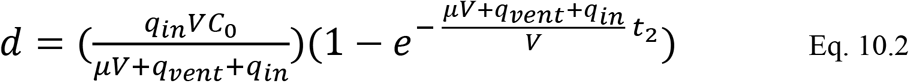

where *t_2_* = receptor’s dwell time (min), and *d* = SARS-CoV-2 infectious dose (PFU).

### SARS-CoV-2 dose-response models

The risk assessment was conducted by following the QMRA framework. Given the lack of dose-response information for SARS-CoV-2, the SARS-CoV data sets (Watanabe, Bartrand, et al., 2010) that had been utilized in various SARS-CoV-2 QMRA studies (Murakami, Miura, et al., 2021; Dada and Gyawali, 2021; Zaneti, Girardi, et al., 2021; Cortellessa, Stabile, et al., 2021) were applied. The risk of infection followed the exponential model (Eq. 11):

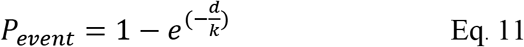

where *P_event_* is the probability of infection per event (probability), and *k* is the optimal dose response function value of 4.1 × 10^2^, which is equivalent to the chance that a single pathogen would initiate an infection response (Watanabe, Bartrand, et al., 2010).

### Risk characterization and sensitivity analysis

To estimate the *P_event_* for a receptor exposed to SARS-CoV-2, the data from the previous steps were integrated into Monte Carlo simulations (MCs) with 10,000 iterations for each condition using Oracle Crystal Ball software version 11.1.2.4.850. MCs is a randomization technique that uses repeated random sampling from distributions given to key input variables in a model, including corresponding uncertainty profiles. The risk of infection was displayed in the 2.5th percentile, mean, and 97.5th percentile using a forest plot in GraphPad Prism version 7.0. It is becoming increasingly important to fully consider the uncertainties, including those in the 97.5th percentile, to maintain a sufficient safety margin for decision-making during the COVID-19 pandemic (Zhang, Ji, et al., 2021). The estimated risk was compared to a benchmark of 1 infection per 20,000 exposed people per event (5 × 10^−5^) (Murakami, Miura, et al., 2021). A sensitivity analysis was also conducted to determine the effects of the input variables on the risk calculation.

### Risk management evaluation

Two risk mitigation interventions were investigated: face mask wearing and ventilation. The universal wearing of face masks has been recommended as a low-cost and efficient means of mitigating virus transmission (WHO, 2020b). Among the different types of face masks, predominantly N95 respirator and surgical and fabric masks are used worldwide. Viral filtration efficiency (VFE) characterized using a bacteriophage MS2 following the ASTM F2101-14 standard testing method has revealed 99.8%–100% VFE for N95 respirators, 99.3%–99.8% VFE for surgical masks, and 54.8%–92.1% for denim fabric masks (Whiley, Keerthirathne, et al., 2020) (Table S1). MS2 bacteriophages were selected as the model microbes because they are two to three times smaller in size than SARS-COV-2 (70–90 nm in diameter).

Ventilation is also an important element used to control indoor air quality in public toilets. Depending on the applicable regulatory building standard, either the installation of a mechanical ventilation system or the use of natural ventilation may be necessary. The effect of air change rates on SARS-CoV-2 transmission risk was considered in this study. The DIN 1946 ventilation standard is generally applied in public toilets. For the pandemic, the WHO has also suggested that ventilation in indoor spaces with aerosol-generating potential should be greater than or equal to 12 ACH (WHO, 2021). In this study, five air change rates were tested: 0 ACH (no ventilation), 0.5 ACH (poor ventilation), 10 ACH (DIN 1946 ventilation standard for public toilets), 12 ACH (WHO-recommended standard ventilation), and 20 ACH (extreme ventilation) (Table S1).

## RESULTS AND DISCUSSION

### Infection risk from respiratory transmission in public toilets

The risk of infection from SARS-CoV-2 transmission through three respiratory exposure scenarios, namely, breathing, breathing with a cough, and breathing with a sneeze, were characterized in this study. The probability of infection per event was not found to be significantly different between men and women (p > 0.05; Mann–Whitney *U* test) across all scenarios (Figure 2 and Table S2). Although men usually have a higher breathing rate (Brochu et al., 2006), which results in greater exposure to viruses, men spend on average around 22% less time in toilets than women (Gwynne, Hunt, et al., 2019), leading to a reduced risk of virus transmission. When a receptor without a protective mask was in an unventilated public toilet, the infection risk (97.5^th^ percentile) was 9.87 × 10^−4^ for men and 1.17 × 10^−3^ for women in the infector breathing scenario. Interestingly, the risk values increased sharply when additional viral loads were expelled into the air by an infector either sneezing or coughing (Figure 2 and Table S2). Coughing and sneezing can produce saliva droplets of various sizes (Duguid, 1946; Han, Weng, et al., 2013; Loudon and Roberts, 1967) and thus generate infectious virus-containing aerosols in public toilet facilities. For breathing with a cough, the 97.5th percentile of risk was 2.17 × 10^−1^ for men and 2.15 × 10^−1^ for women. Similarly, sneezing increased the risk of infection to 3.66 × 10^−1^ and 3.67 × 10^−1^ for men and women, respectively. All the scenarios demonstrated higher risks than the 5 × 10^−5^ benchmark value (Murakami, Miura, et al., 2021). We therefore showed that receptors had a high risk of infection when using an unventilated public toilet without wearing a protective mask.

**Figure 2.**
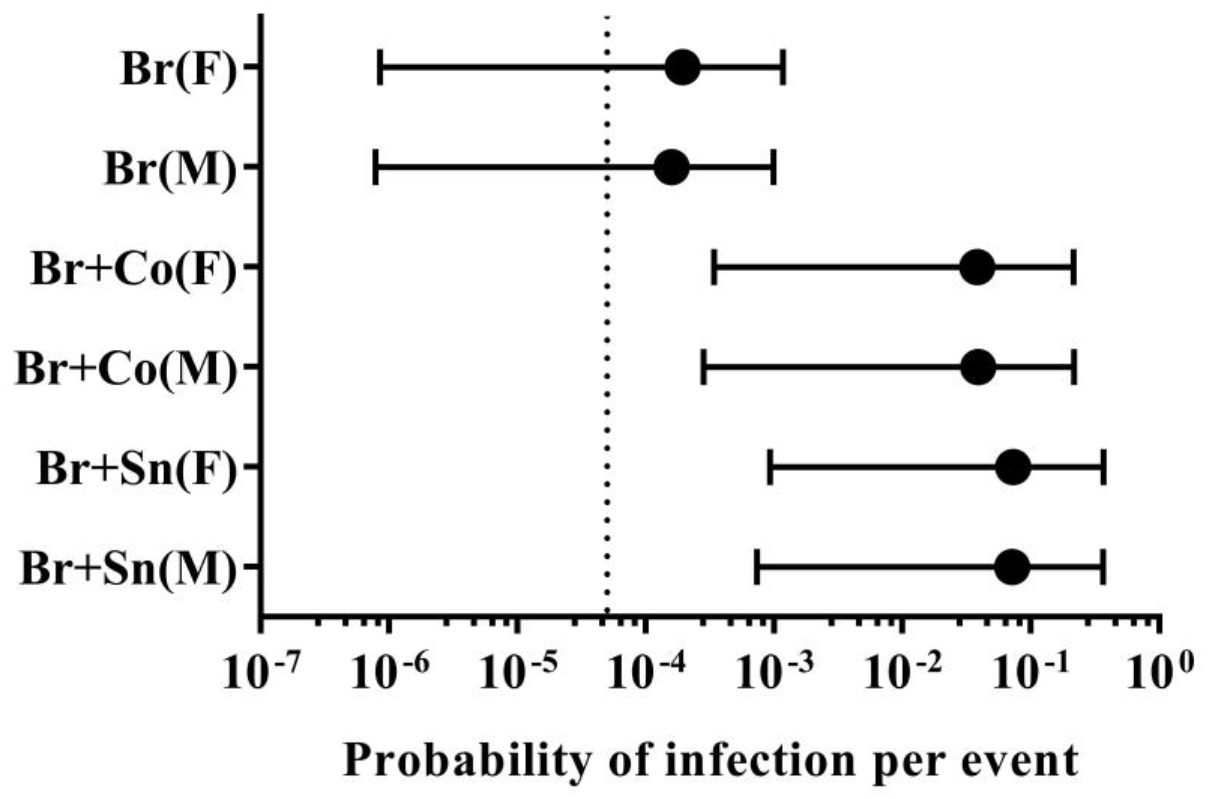
Risk of infection per event for a male (M) or female (F) receptor in three virus-generating scenarios: an infector breathing (Br), breathing with a cough (Br+Co), and breathing with a sneeze (Br+Sn). The forest plots show the mean values in solid circles and 95% confidence intervals (ranging from the 2.5th [left whiskers] to the 97.5th [right whiskers] percentiles). The dashed line indicates the 5 × 10^−5^ benchmark value.

### Risk mitigation: face mask wearing

#### Face mask wearing in either an infector or a receptor

Because it delivered the highest risk, the scenario with an infector breathing with a sneeze was selected to further evaluate the effectiveness of face mask wearing to reduce infection risk in a receptor. The probabilities of infection per event for different mask types are shown in Figure 3 and Table S3. All types of face masks considerably reduced the risk of infection. For example, an N95 respirator could lead to an approximately 2-log reduction when worn by an infector and a 3-log reduction when worn by a receptor. Interestingly, mask wearing by a receptor reduced the risk of transmission to lower levels than when a mask was worn by an infector. However, the results indicated that face mask wearing either by an infector or a receptor could still not decrease the risk to below the suggested 5 × 10^−5^ benchmark. A high risk of virus transmission in confined spaces like public toilets is therefore still possible even if an N95 respirator or surgical mask is worn. This could be because the risk of infection is associated with several factors, including a high concentration of virus aerosols in the ambient air due to insufficient ventilation, the long dwell times of toilet users, the low inactivation rate of SAR-CoV-2, and the inadequate efficiency of protective masks (Stabile, Pacitto, et al., 2021; Gwynne, Hunt, et al., 2019; Schuit, Ratnesar-Shumate, et al., 2020; Whiley, Keerthirathne, et al., 2020). We also observed an at least 90% (1-log) reduction in risk when an infector wore a denim fabric mask.

**Figure 3.**
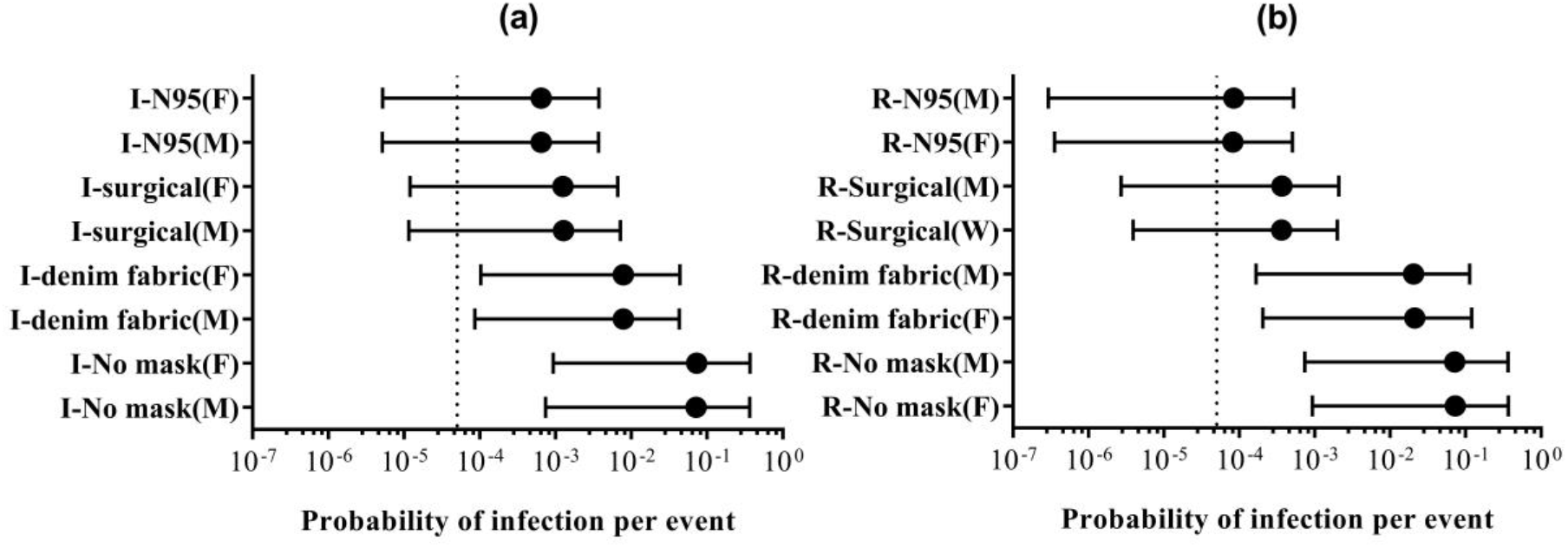
Risk of infection per event among male (M) and female (F) receptors in the scenario with an infector (I) breathing with a sneeze (Br+Sn) when (a) only the infector wore different types of masks and (b) only the receptor (R) wore different types of masks. The forest plots show the mean values in solid circles and 95% confidence intervals (ranging from the 2.5th [left whiskers] to the 97.5th [right whiskers] percentiles). The dashed line indicates the 5 × 10^−5^ benchmark value.

#### Face mask wearing in both an infector and a receptor

In a scenario with an infector breathing with a sneeze, the infection risk could be further reduced if both the infector and receptor wear masks (Table S3). The receptor’s gender did not affect the receptor’s risk much in any of the conditions. The risks to a female receptor are illustrated in Figure 4. When a receptor wore an N95 respirator, the 97.5th percentile infection risk was reduced to below the 5 × 10^−5^ benchmark no matter the type of mask, whether an N95 respirator or a surgical or denim fabric mask, worn by the infector. When a receptor wore a surgical mask, the risk was also reduced to below the benchmark with the exception of the case where the infector wore a fabric mask. Denim fabric masks, on the other hand, may not provide sufficient protection even when worn by both the infector and receptor. This study supports the recommendation for a person, as a receptor, to use a surgical mask or an N95 respirator as personal protective equipment to minimize the associated risk of infection in unventilated public toilets and potentially in other confined communal spaces. In general, wearing a mask was shown to be one of the most low-cost, simple yet effective intervention measures to minimize transmission risk, which is consistent with reports from other studies (Asadi, Cappa, et al., 2020; Chu, Akl, et al., 2020; WHO, 2020b; Cheng, Cheng, et al., 2021; Goyal, Reeves, et al., 2021).

**Figure 4.**
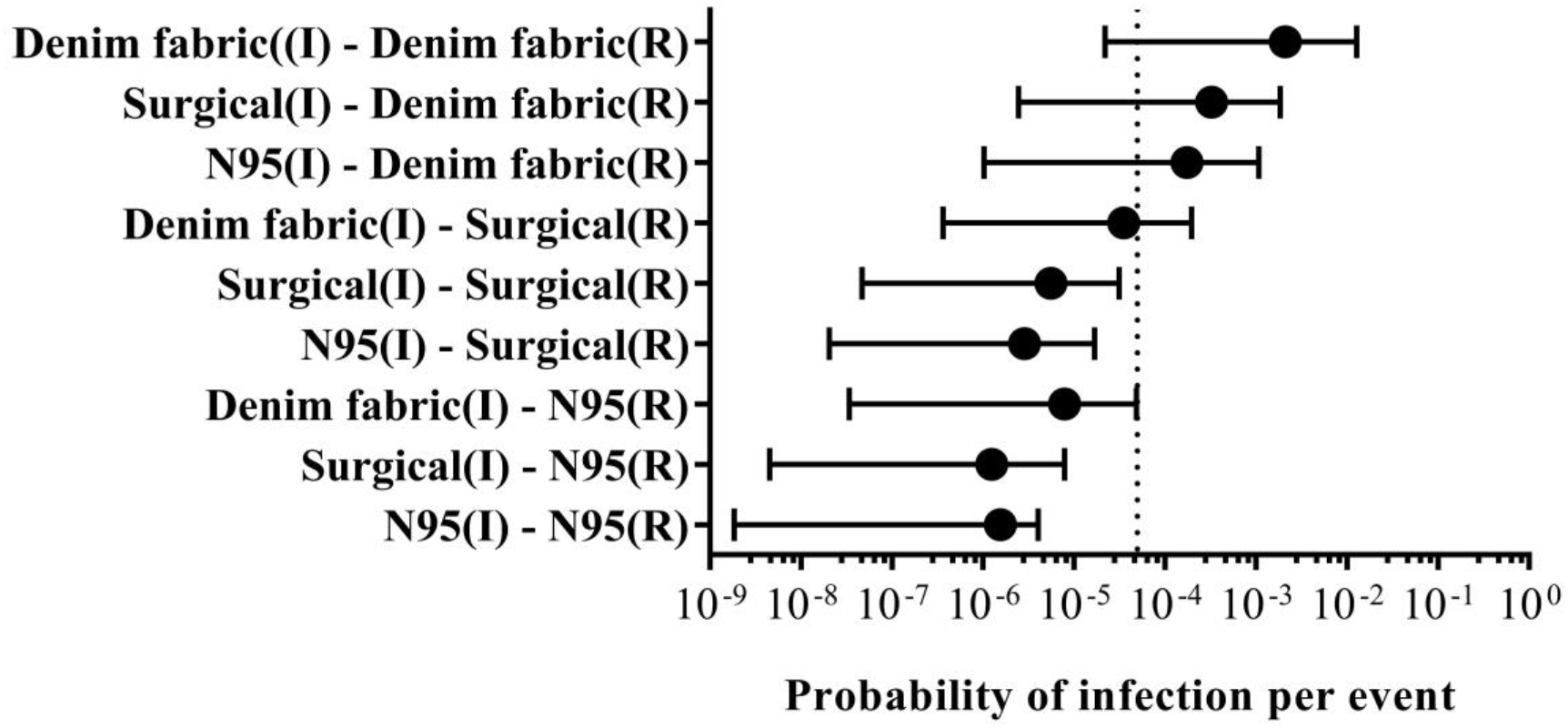
The risk of infection per event in a scenario with an infector (I) breathing with a sneeze (Br+Sn) when both the infector and receptor (R) wore different types of masks. The risks to a female receptor are represented because no gender effect was evident. The forest plots show the mean values in solid circles and 95% confidence intervals (ranging from the 2.5th [left whiskers] to the 97.5th [right whiskers] percentiles). The dashed line indicates the 5 × 10^−5^ benchmark value.

### Risk mitigation: ventilation

#### Single measure: ventilation

The effects of ventilation (0–20 ACH) were characterized for three virus-generating scenarios: infector breathing, breathing with a cough, and breathing with a sneeze (Table S4). The breathing with a sneeze scenario delivered the highest risk, and this worst-case condition was therefore further assessed to determine the effects of ventilation using a representative female receptor (Figure 5). The results showed that increasing ACH did not significantly mitigate the risk of COVID-19 infection in the public toilet setting. Even at the 12 ACH suggested by the WHO (2021) and the extreme condition of 20 ACH, the 97.5th percentile probabilities of infection per event for the female receptor were still at the levels of 2.52 × 10^−1^ and 2.10 × 10^−1^, respectively. Although a high ventilation rate has been suggested as a way to reduce the number of virus-containing droplets and aerosols in the air (WHO, 2021; Li, Qian, et al., 2021; Morawska, Tang, et al., 2020; Stabile, Pacitto, et al., 2021), the continuous expelling of the SARS-CoV-2 virus from breathing and/or sneezing by an infector without a mask appeared to be a significant cause of virus aerosol accumulation in ambient air. This study demonstrated that indoor ventilation alone cannot effectively reduce SARS-CoV-2 transmission risk in a public toilet setting and is less effective in risk reduction than face mask wearing. Another study similarly found that masks could reduce the infection risk caused by the Middle Eastern respiratory syndrome coronavirus in an indoor hospital setting better than ventilation (Adhikari, Chabrelie, et al., 2019).

**Figure 5.**
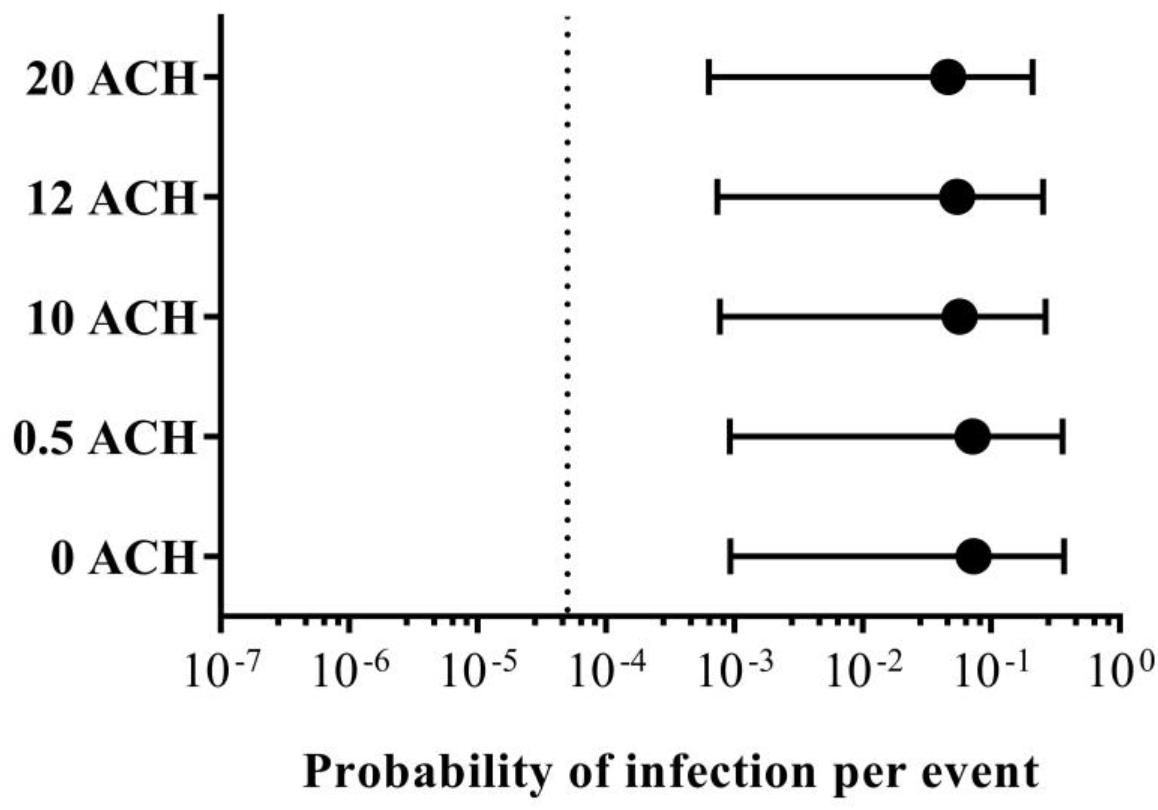
The risk of infection per event in a scenario with an infector breathing with a sneeze (Br+Sn) with 0, 0.5, 10, 12, and 20 air changes per hour (ACH). The risks to a female receptor are represented because no gender effect was evident. The forest plots show the mean values in solid circles and 95% confidence intervals (ranging from the 2.5th [left whiskers] to the 97.5th [right whiskers] percentiles). The dashed line indicates the 5 × 10^−5^ benchmark value.

#### Double measure: ventilation and face mask wearing

The additional measure of mask wearing was further investigated for its combined effectiveness in mitigating SARS-CoV-2 transmission risk when used simultaneously with increased ventilation. In the virus-generating scenario with an infector breathing with a sneeze, mask wearing by both the infector and receptor was assessed using ventilation of 10, 12, and 20 ACH (Figure 6). When compared with no ACH (Figure 4), ventilation across all ACH values did not reduce the infection risk to below the benchmark in any of the following four cases: denim fabric mask wearing by the receptor and all three types of masks worn separately by the infector, and surgical mask wearing by the receptor and denim fabric mask wearing by the infector. Consequently, we reiterate that ventilation did not impact the risk mitigation for SARS-CoV-2 transmission in a public toilet setting, especially in confined toilet cubicle conditions. Face mask wearing should therefore be promoted as a normal practice when entering public indoor spaces.

**Figure 6.**
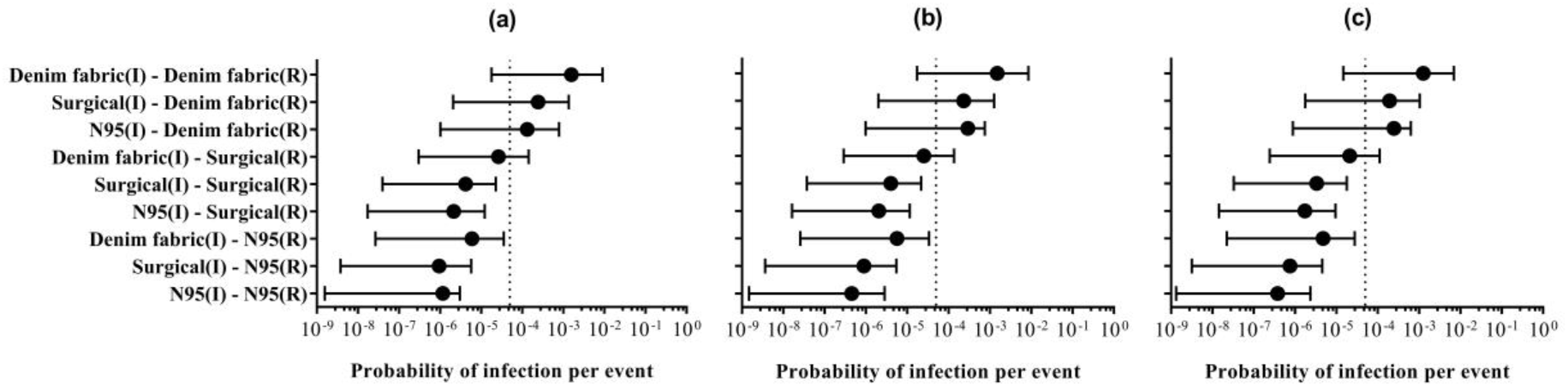
The risk of infection per event in a scenario with an infector (I) breathing with a sneeze (Br+Sn) when the infector wore different types of masks, the receptor (R) wore different types of masks, and the air change per hour was (a) 10 ACH, (b) 12 ACH, and (c) 20 ACH. The risks to a female receptor are represented because no gender effect was evident. The forest plots show the mean values in solid circles and 95% confidence intervals (ranging from the 2.5th [left whiskers] to the 97.5th [right whiskers] percentiles). The dashed line indicates the 5 × 10^−5^ benchmark value.

### Sensitivity analysis of input parameters

A sensitivity analysis of the QMRA was conducted to identify the input variables that most contributed to the risk estimation. For all three transmission scenarios, namely, an infector breathing (Br), breathing with a cough (Br+Co), and breathing with a sneeze (Br+Sn), the concentration of SARS-CoV-2 virus in gc/μL droplets (saliva and mucus) was the most sensitive parameter, accounting for 34.3%–42.9% of the uncertainty in the probability of infection transmission to either male or female receptors (Figure 7 and Table S5). The second and third most sensitive parameters were the infector’s expelled volume and the receptor’s dwell time, respectively. Since breathing with a sneeze was the highest virus-generating risk scenario, a sensitivity analysis was performed in which both the infector and receptor wore masks (Table S6). Virus concentrations in gc/μL, sneeze volume, and the receptor’s dwell time were the three parameters that most influenced infection risk. Since controlling for virus concentrations and an infector’s expelled volume are a challenge, particularly among asymptomatic patients, individuals should avoid spending prolonged time in closed indoor settings (Dancer, Li, et al., 2021; Stabile, Pacitto, et al., 2021).

**Figure 7.**
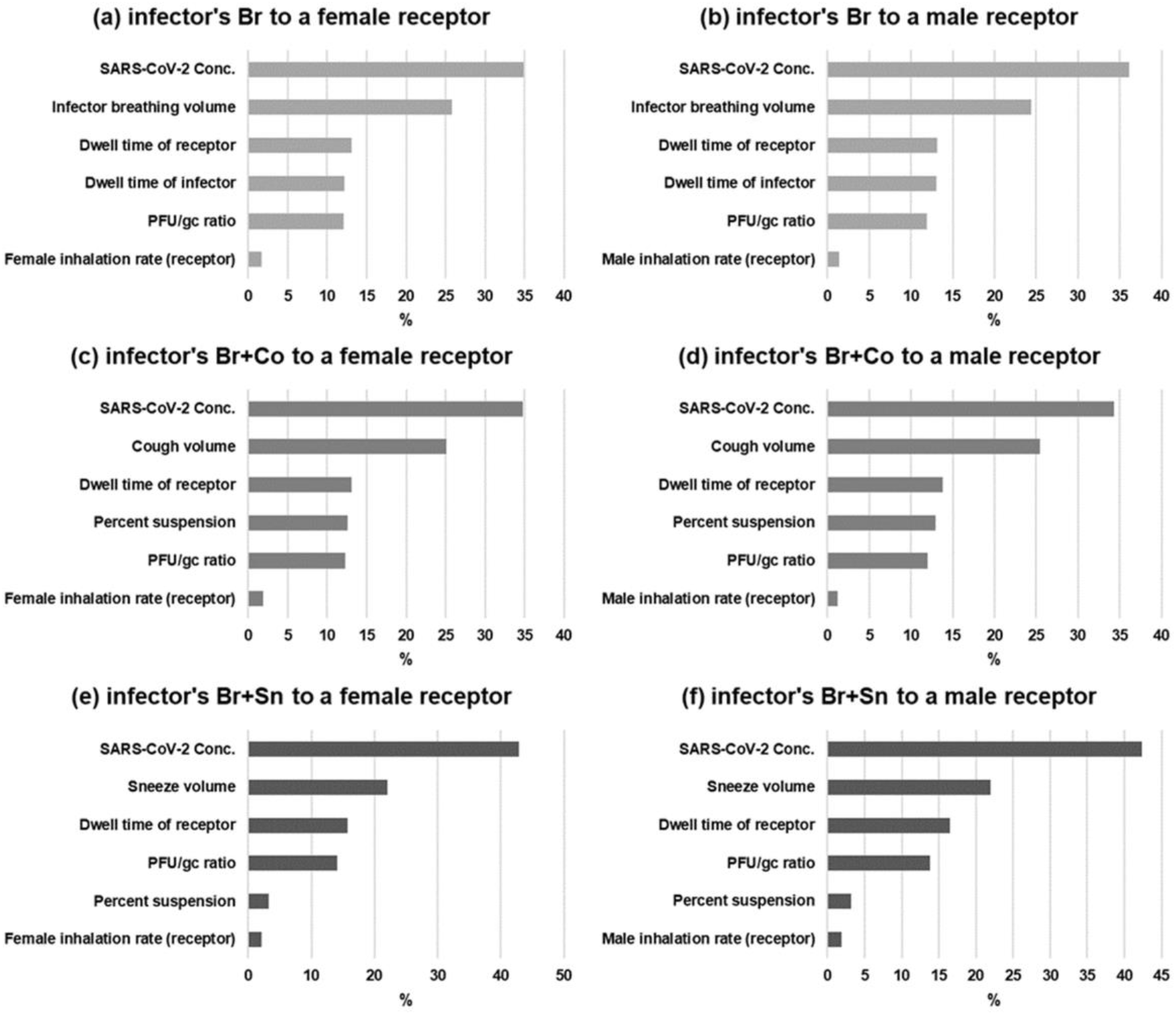
The sensitivity analysis representing the contribution of input variables to the risk of infection per event for male and female receptors in three transmission scenarios, namely, with the infector breathing (Br), breathing with a cough (Br+Co), and breathing with a sneeze (Br+Sn): (a) infector’s Br to a female receptor, (b) infector’s Br to a male receptor, (c) infector’s Br+Co to a female receptor, (d) infector’s Br+Co to a male receptor, (e) infector’s Br+Sn to a female receptor, and (f) infector’s Br+Sn to a male receptor.

### Limitations of this study and future perspectives

While this study evaluated the risk of SARS-CoV-2 transmission according to the QMRA framework, its limitations and uncertainties should be carefully acknowledged. SARS-CoV-2 concentrations in gc/μL, the most sensitive parameter affecting the calculation of risk, are subject to natural variations in the saliva and mucus of infected patients (Azzi, Carcano, et al., 2020; Wölfel, Corman, et al., 2020). In this study, 251 swab test *Ct* values were used to represent the virus levels in Thai patients. Due to the lack of a standard curve from Thai hospital laboratories, we used a published standard curve of the N2 gene (Sherchan, Shahin, et al., 2020) to estimate the virus concentrations in this study. However, heterogeneity in published standard curves for SARS-CoV-2 has been observed (Bivins, Kaya, et al., 2021). Moreover, variations in technical and laboratory analyses (e.g., data analysis methods and control materials) could intensify biases, leading to variability in the calculated virus concentrations (Bivins, Kaya, et al., 2021; Kongprajug, Chyerochana, et al., 2020). Adhering to standards and quality control measures is therefore underlined in order to support data sharing and referencing for future research, especially for emerging infectious diseases. However, even with consideration of the uncertainties mentioned above, the calculated virus concentrations in mucus used in this study, which ranged from 4.4 × 10^−1^ to 6.4 × 10^8^ gc/μL, were in agreement with those from another report (Schijven, Vermeulen, et al., 2021).

We chose to evaluate three virus-generating scenarios: an inceptor breathing, breathing with a cough, and breathing with a sneeze. However, infectors may sneeze and/or cough more than once depending on the individuals’ symptoms. Coughing is the predominant symptom in COVID-19 (Wang, Yang, et al., 2020), rendering coughing potentially more important than sneezing. Nevertheless, some studies have suggested that airborne transmission of infectious diseases is possible without coughing or sneezing and simply from exhaled breath from individuals who show barely any symptoms (Asadi, Bouvier, et al., 2020). In addition, the lack of dose-response information and risk of infection benchmarks for SARS-CoV-2 poses a challenge when evaluating its infection risk. We assumed that the dose-response of SARS-CoV-2 was similar to that indicated in the SARS-CoV data (Watanabe, Bartrand, et al., 2010), which has been utilized in various QMRA studies of SARS-CoV-2 (Murakami, Miura, et al., 2021; Dada and Gyawali, 2021; Zaneti, Girardi, et al., 2021; Cortellessa, Stabile, et al., 2021). With the recent emergence of various SARS-CoV-2 variants, much remains unknown regarding the behavior and characteristics of this virus. This study used the available inactivation coefficients of SARS-CoV-2 at 20°C (Schuit, Ratnesar-Shumate, et al., 2020), which could have overestimated the calculated risks in Thailand given its average daily temperature of 27.48°C (Denpetkul & Phosri, 2021).

Furthermore, the scope of this study excluded the risk of SARS-CoV-2 respiratory transmission potentially produced by toilet flushing, as well as other transmission risks (e.g., direct splashing and surface transmission). By integrating all the known risk sources, comprehensive knowledge regarding risk estimation could be achieved to accurately inform public health policy and further help reduce transmission risk. It is apparent that research related to SARS-CoV-2 is continuing, and additional data will greatly benefit future studies aiming to better understand its characteristics. The QMRA-based risk models developed in this study could facilitate future risk assessments through modifications for particular risk scenarios and the updating of the input parameters based on newly available data. Such improved risk models will be crucial tools in assessing the impact of different risk mitigation strategies during the COVID-19 and future pandemics.

## CONCLUSIONS

Indoor public toilet facilities could be hubs of virus transmission during the COVID-19 pandemic. This study investigated the risk of airborne transmission of SARS-CoV-2 in public toilets for three virus-generating scenarios: an infector breathing, breathing with a cough, and breathing with a sneeze. The risk analysis, which followed the QMRA framework, revealed that the highest risk was when an asymptomatic or symptomatic infector sneezed. Both genders were found to be exposed to similar risks. Toilet ventilation systems cannot effectively mitigate transmission risk, so an effective intervention would be for public toilet users to wear either surgical masks or N95 respirators.

## Acknowledgments

This work was financially supported by the Chulabhorn Research Institute (grant no. 312/3057) and the Mahidol University. The authors would like to acknowledge Associate Professor Pornsawan Leaungwutiwong from the Department of Microbiology and Immunology, Faculty of Tropical Medicine, Mahidol University for providing the COVID-19 Ct values from swab testing.

## Supplementary Material

Supplementary material for this manuscript is available.

