## Supplemental tables for "Effects of face masks and ventilation on the risk of SARS-CoV-2 respiratory transmission in public toilets: a quantitative microbial risk assessment"

[1]Faculty of Tropical Medicine, Mahidol University, Bangkok, Thailand 10400; [2]Environmental Engineering and Disaster Management Program, School of Multidisciplinary, Mahidol University, Kanchanaburi Campus, Sai Yok, Kanchanaburi, Thailand, 71150; [3]Research Laboratory of Biotechnology, Chulabhorn Research Institute, Bangkok, Thailand 10210; [4]Center of Excellence on Environmental Health and Toxicology (EHT), Ministry of Education, Bangkok, Thailand 10400

*

**Table S1** Probability density functions of the model input data for the quantitative microbial risk assessment simulation

| Parameter | Unit | Description | Distribution | Sources |
|---|---|---|---|---|
| *A* | gc/μL | Concentration of SARS-CoV-2 in mucus | Triangular (0.44, 4.94, $6.45 \times 10^8$) | This study, Sherchan et al., 2020 |
| *R* | PFU/gc | Conversion ratio of PFU to gc | Uniform (0.001, 0.01) | Pitol & Julian, 2021 |
| $q_{br}$ | pL/min | Volumetric flow of droplets from an infector's exhalation | Uniform (0.005, 6) | Schijven et al., 2021 |
| $V_{co}$ | pL/cough | Volume of aerosol expelled per cough | Uniform (9.5, $1.3 \times 10^4$) | Schijven et al., 2021 |
| $V_{sn}$ | pL/sneeze | Volume of aerosol expelled per sneeze | Uniform ($2.1 \times 10^3$, $8 \times 10^4$) | Schijven et al., 2021 |
| *V* | L | Volume of air in a cubicle | Point (3240) | This study |
| *F* | Dimensionless | Volumetric ratio of aerosols suspended per total droplets expelled | For breathing: point (1) | Fabian et al., 2011 |
| | | | For coughing: uniform (0.04, 0.49) | Duguid, 1946; Han et al., 2013; Loudon & Roberts, 1967 |
| | | | For sneezing: uniform (0.04, 0.1224) | Duguid, 1946; Han et al., 2013 |
| $t_1$ | min | An infector's dwell time in the toilet cubicle | Men: log-normal (2.22, 1.53) | Gwynne et al., 2019 |
| | | | Women: log-normal (2.78, 1.80) | |
| $t_2$ | min | A receptor's dwell time in the toilet cubicle | Men log-normal (2.22, 1.53) | Gwynne et al., 2019 |
| | | | Women lognormal (2.78, 1.80) | |
| $q_{in}$ | L/min | Inhalation rate | Men uniform (8.36, 19.74) | Brochu et al., 2006 |
| | | | Women uniform (6.40, 13.78) | |
| $\mu$ | $min^{-1}$ | Inactivation rate in air (20%–70% relative humidity levels) | Point (0.008) | Schuit et al., 2020 |
| N95 respirator | Removal ratio | 99.8%–100% VFE | Uniform (0.000, 0.002) | Whiley et al., 2020 |

| | | | | |
|---|---|---|---|---|
| Surgical mask | Removal ratio | 99.3%–99.8% VFE | Uniform (0.002, 0.007) | Whiley et al., 2020 |
| Fabric mask | Removal ratio | 54.8%–92.1% VFE | Uniform (0.079, 0.452) | Whiley et al., 2020 |
| ACH | $hr^{-1}$ | ACH | No ventilation<br>ACH = 0 | |
| | | | Poor ventilation<br>ACH = 0.5 | |
| | | | Standard ventilation<br>ACH = 10 | Ventilation standard for public toilet DIN1946 |
| | | | Recommended ventilation<br>ACH = 12 (with aerosol-generating potential) | WHO, 2021 |
| | | | Extreme ventilation<br>ACH =20 (operating rooms) | |

ACH, air changes per hour; PFU, plaque-forming unit; VFE, viral filtration efficiency

**Table S2** Probability of SARS-CoV-2 infection per event for three scenarios (with an infector breathing, breathing with a cough, and breathing with a sneeze) where the infector and receptor are not wearing masks and without ventilation

| | Risk value | | | | | | | | |
|---|---|---|---|---|---|---|---|---|---|
| **Descriptive statistics**[a] | **Breathing** | | | **Breathing + a cough** | | | **Breathing + a sneeze** | | |
| | **Men** | **Women** | ***p*-value**[b] | **Men** | **Women** | ***p*-value** | **Men** | **Women** | ***p*-value** |
| P2.5 | 7.88E-07 | 8.55E-07 | 0.645 | 2.84E-04 | 3.41E-04 | 0.959 | 7.40E-04 | 9.29E-04 | 0.742 |
| Mean | 1.60E-04 | 1.94E-04 | | 3.91E-02 | 3.84E-02 | | 7.16E-02 | 7.26E-02 | |
| P97.5 | 9.87E-04 | 1.17E-03 | | 2.17E-01 | 2.15E-01 | | 3.66E-01 | 3.67E-01 | |

[a] P2.5 and P97.5 refer to the 2.5th and 97.5th percentile values, respectively

[b] Mann–Whitney *U* test for the non-normal data between the male and female risk

**Table S3** Probability of SARS-CoV-2 infection per event in the scenario with an infector breathing with a sneeze in an unventilated public toilet when either the infector or receptor or both are wearing different mask types

| Infector wearing an N95 respirator | Receptor mask types | | | | | | | |
|---|---|---|---|---|---|---|---|---|
| | No mask | | N95 respirator | | Surgical mask | | Denim fabric mask | |
| Descriptive statistics[a] | Men | Women | Men | Women | Men | Women | Men | Women |
| P2.5 | 5.16E-06 | 5.20E-06 | 1.85E-09 | 1.85E-09 | 2.18E-08 | 2.05E-08 | 1.16E-06 | 1.03E-06 |
| Mean | 6.43E-04 | 6.43E-04 | 6.47E-07 | 6.27E-07 | 2.97E-06 | 2.86E-06 | 1.68E-04 | 1.75E-04 |
| P97.5 | 3.68E-03 | 3.72E-03 | 4.03E-06 | 4.06E-06 | 1.75E-05 | 1.69E-05 | 1.04E-03 | 1.08E-03 |
| **Infector wearing a surgical mask** | **Receptor mask types** | | | | | | | |
| | **No mask** | | **N95 respirator** | | **Surgical mask** | | **Denim fabric mask** | |
| **Descriptive statistics** | **Men** | **Women** | **Men** | **Women** | **Men** | **Women** | **Men** | **Women** |
| P2.5 | 1.16E-05 | 1.20E-05 | 3.98E-09 | 4.55E-09 | 4.48E-08 | 4.69E-08 | 2.52E-06 | 2.46E-06 |
| Mean | 1.27E-03 | 1.25E-03 | 1.27E-06 | 1.25E-06 | 5.71E-06 | 5.56E-06 | 3.25E-04 | 3.24E-04 |
| P97.5 | 7.17E-03 | 6.64E-03 | 7.72E-06 | 7.92E-06 | 3.13E-05 | 3.15E-05 | 1.88E-03 | 1.85E-03 |
| **Infector wearing a denim fabric mask** | **Receptor mask types** | | | | | | | |
| | **No mask** | | **N95 respirator** | | **Surgical mask** | | **Denim fabric mask** | |
| **Descriptive statistics** | **Men** | **Women** | **Men** | **Women** | **Men** | **Women** | **Men** | **Women** |
| P2.5 | 8.56E-05 | 1.03E-04 | 3.81E-08 | 3.40E-08 | 3.49E-07 | 3.67E-07 | 1.92E-05 | 2.19E-05 |
| Mean | 7.84E-03 | 7.83E-03 | 7.91E-06 | 7.94E-06 | 3.56E-05 | 3.52E-05 | 2.11E-03 | 2.10E-03 |
| P97.5 | 4.29E-02 | 4.39E-02 | 4.96E-05 | 4.89E-05 | 2.03E-04 | 1.97E-04 | 1.30E-02 | 1.28E-02 |
| **Infector wearing no mask** | **Receptor mask types** | | | | | | | |
| | **No mask** | | **N95 respirator** | | **Surgical mask** | | **Denim fabric mask** | |
| **Descriptive statistics** | **Men** | **Women** | **Men** | **Women** | **Men** | **Women** | **Men** | **Women** |
| P2.5 | 7.40E-04 | 9.29E-04 | 2.94E-07 | 3.56E-07 | 2.72E-06 | 3.91E-06 | 1.67E-04 | 2.04E-04 |
| Mean | 7.16E-02 | 7.26E-02 | 8.41E-05 | 8.19E-05 | 3.66E-04 | 3.64E-04 | 2.05E-02 | 2.12E-02 |
| P97.5 | 3.66E-01 | 3.67E-01 | 5.26E-04 | 5.04E-04 | 2.08E-03 | 1.99E-03 | 1.13E-01 | 1.20E-01 |

[a] P2.5 and P97.5 refer to the 2.5th and 97.5th percentile values, respectively

**Table S4** Probability of SARS-CoV-2 infection per event for three scenarios (an infector breathing, breathing with a cough, and breathing with a sneeze) where the infector and receptor are not wearing masks but with different air change rates

| Breathing | | | | | | | | | | |
|---|---|---|---|---|---|---|---|---|---|---|
| **Descriptive statistics[a]** | **0 ACH** | | **0.5 ACH** | | **10 ACH** | | **12 ACH** | | **20 ACH** | |
| | **Men** | **Women** | **Men** | **Women** | **Men** | **Women** | **Men** | **Women** | **Men** | **Women** |
| **P2.5** | 7.88E-07 | 8.55E-07 | 7.80E-07 | 8.38E-07 | 6.01E-07 | 5.83E-07 | 5.67E-07 | 5.48E-07 | 4.55E-07 | 4.31E-07 |
| **Mean** | 1.60E-04 | 1.94E-04 | 1.56E-04 | 1.88E-04 | 9.90E-05 | 1.11E-04 | 9.09E-05 | 1.00E-04 | 6.66E-05 | 7.02E-05 |
| **P97.5** | 9.87E-04 | 1.17E-03 | 9.62E-04 | 1.14E-03 | 5.74E-04 | 6.15E-04 | 5.21E-04 | 5.42E-04 | 3.65E-04 | 3.59E-04 |
| **Breathing with a cough** | | | | | | | | | | |
| **Descriptive statistics[a]** | **0 ACH** | | **0.5 ACH** | | **10 ACH** | | **12 ACH** | | **20 ACH** | |
| | **Men** | **Women** | **Men** | **Women** | **Men** | **Women** | **Men** | **Women** | **Men** | **Women** |
| **P2.5** | 2.84E-04 | 3.41E-04 | 2.82E-04 | 3.39E-04 | 2.45E-04 | 2.77E-04 | 2.42E-04 | 2.65E-04 | 2.12E-04 | 2.26E-04 |
| **Mean** | 3.91E-02 | 3.84E-02 | 3.86E-02 | 3.79E-02 | 3.16E-02 | 2.97E-02 | 3.04E-02 | 2.84E-02 | 2.63E-02 | 2.39E-02 |
| **P97.5** | 2.17E-01 | 2.15E-01 | 2.14E-01 | 2.11E-01 | 1.70E-01 | 1.64E-01 | 1.63E-01 | 1.56E-01 | 1.41E-01 | 1.30E-01 |
| **Breathing with a sneeze** | | | | | | | | | | |
| **Descriptive statistics[a]** | **0 ACH** | | **0.5 ACH** | | **10 ACH** | | **12 ACH** | | **20 ACH** | |
| | **Men** | **Women** | **Men** | **Women** | **Men** | **Women** | **Men** | **Women** | **Men** | **Women** |
| **P2.5** | 7.40E-04 | 9.29E-04 | 7.33E-04 | 9.25E-04 | 6.03E-04 | 7.68E-04 | 5.84E-04 | 7.39E-04 | 5.11E-04 | 6.34E-04 |
| **Mean** | 7.16E-02 | 7.26E-02 | 7.09E-02 | 7.16E-02 | 5.87E-02 | 5.68E-02 | 5.66E-02 | 5.43E-02 | 4.93E-02 | 4.60E-02 |
| **P97.5** | 3.66E-01 | 3.67E-01 | 3.61E-01 | 3.60E-01 | 2.91E-01 | 2.65E-01 | 2.76E-01 | 2.52E-01 | 2.34E-01 | 2.10E-01 |

ACH, air changes per hour

[a] P2.5 and P97.5 refer to the 2.5th and 97.5th percentile values, respectively

**Table S5** Sensitivity analysis of SARS-CoV-2 infection per event for three scenarios (an infector breathing, breathing with a cough, and breathing with a sneeze) without ventilation and where the infector and receptor are not wearing masks

| **Breathing** | | | **Breathing + a cough** | | | **Breathing + a sneeze** | | |
|---|---|---|---|---|---|---|---|---|
| **Sensitivity parameter** | **Men** | **Women** | **Sensitivity parameter** | **Men** | **Women** | **Sensitivity parameter** | **Men** | **Women** |
| SARS-CoV-2 concentration | 36.1% | 34.9% | SARS-CoV-2 concentration | 34.3% | 34.8% | SARS-CoV-2 concentration | 42.4% | 42.9% |
| Infector breathing volume | 24.4% | 25.8% | Cough volume | 25.4% | 25.1% | Sneeze volume | 22.0% | 22.0% |
| Dwell time of receptor | 13.1% | 13.1% | Dwell time of receptor | 13.8% | 13.1% | Dwell time of receptor | 16.5% | 15.7% |
| Dwell time of infector | 13.0% | 12.2% | Percent suspension | 12.9% | 12.6% | PFU/gc ratio | 13.8% | 14.1% |
| PFU/gc ratio | 11.9% | 12.1% | PFU/gc ratio | 12.0% | 12.3% | Percent suspension | 3.2% | 3.2% |
| Inhalation rate (receptor) | 1.4% | 1.7% | Inhalation rate (receptor) | 1.2% | 1.9% | Inhalation rate (receptor) | 1.9% | 2.1% |
| Other | 0.1% | 0.2% | Other | 0.4% | 0.2% | Other | 0.2% | 0.0% |

gc, gene copies; PFU, plaque-forming unit

**Table S6** Sensitivity analysis of SARS-CoV-2 infection per event with an infector breathing with a sneeze in an unventilated public toilet when both the infector and receptor are wearing different types of masks

| **Infector wearing surgical mask** | **Receptor mask type** | | | |
|---|---|---|---|---|
| | **N95 respirator** | | **Surgical mask** | |
| **Sensitivity parameter** | **Men** | **Women** | **Men** | **Women** |
| SARS-CoV-2 concentration | 30.1% | 30.6% | 38.8% | 37.7% |
| Sneeze volume | 17.6% | 17.0% | 22.9% | 21.1% |
| Dwell time of receptor | 13.9% | 11.8% | 14.1% | 16.4% |
| PFU/gc ratio | 10.6% | 10.4% | 13% | 12.8% |
| Viral penetration of mask (receptor) | 23.4% | 24.2% | 5.4% | 5.5% |
| Percent suspension | 2.7% | 3.1% | 3.3% | 3.3% |
| Inhalation rate (receptor) | 1.0% | 2.4% | 1.7% | 2.4% |
| Viral penetration of mask (infector) | 0.1% | 0.1% | 0.2% | 0.2% |
| Other | 0.6% | 0.4% | 0.6% | 0.6% |
| **Infector wearing N95 mask** | **Receptor mask type** | | | |
| | **N95 respirator** | | **Surgical mask** | |
| **Sensitivity parameter** | **Men** | **Women** | **Men** | **Women** |
| SARS-CoV-2 concentration | 30.8% | 30.9% | 37.4% | 37.4% |
| Sneeze volume | 16.8% | 17.1% | 20.4% | 20.6% |
| Dwell time of receptor | 11.6% | 9.8% | 14.0% | 12.4% |
| PFU/gc ratio | 10.8% | 11.4% | 12.9% | 13.2% |
| Viral penetration of mask (receptor) | 20.7% | 21.5% | 5.6% | 6.1% |
| Viral penetration of mask (infector) | 4.6% | 5.0% | 4.7% | 4.7% |
| Percent of suspension | 2.8% | 2.7% | 3.2% | 3.2% |
| Inhalation rate (receptor) | 1.6% | 1.2% | 1.5% | 2.1% |
| Other | 0.3% | 0.4% | 0.3% | 0.3% |

gc, gene copies; PFU, plaque-forming unit
